# Begging costs rather than food received cause brood size effect on growth in zebra finches

**DOI:** 10.1101/2025.01.29.635459

**Authors:** Marianthi Tangili, Michael Briga, Simon Verhulst

## Abstract

Altricial species rely on parental provisioning for early-life sustenance, and a larger brood size leads to higher levels of competition between siblings for parental resources. Early-life stress can have severe and lifelong effects on Darwinian fitness. Indeed, it is well established that being reared in a larger brood impairs growth and fitness prospects of birds, but the mechanistic underpinnings of this effect are still largely unknown. Specifically, it is not well known to what extent the reduced growth and fitness prospects of nestlings reared in large broods is due to increased resource allocation to competition versus a *per capita* reduction in parental provisioning rate, or a combination of the two. We cross-fostered zebra finch (*Taeniopygia castanotis*) chicks into small and large broods, and recorded their growth as well as the behavior of parents and offspring throughout the nestling period. As in previous experiments, growth rate was higher in small broods. In large broods, chicks begged more and parents invested more time feeding which more than compensated for the difference in brood size. We therefore conclude that the lower growth rate for nestlings raised in large broods must be atleast in part-attributed to increased energy expenditure on begging rather than a reduction in the amount of food received. These results suggest significant energetic costs associated with begging and raise the interesting possibility that brood size would not have negatively affected growth in large broods if chicks had not increased their begging effort due to increased levels of competition in nest.

## Introduction

Early-life stress can have profound and long-lasting effects on the Darwinian fitness of individuals. Animals exposed to adversity during the developmental stages of their lives can exhibit reduced reproductive success (van de Pol et al., 2006) and lifespan (Briga et al., 2017; Tung et al., 2016), altered rates of senescence (Nussey et al., 2007) and differential social behavior (Gerritsma, 2025; Goldenberg & Wittemyer, 2018; Patterson et al., 2022). In birds, early-life stress has been found to affect foraging behavior and mass regulation (Andrews et al., 2015), attractiveness to mates (Holveck & Riebel, 2010; Mainwaring et al., 2012), telomere shortening rates (Boonekamp et al., 2014; Nettle et al., 2015) as well as inflammation (Nettle et al., 2017). Early-life challenges can have direct effects on Darwinian fitness by affecting growth (de Kogel, 1997; Nilsson & Svensson, 2001), recruitment (Schwagmeyer & Mock, 2008), nestling survival (Mock et al., 2009), foraging behavior (Andrews et al., 2015) and reproductive success (Spagopoulou et al., 2020).

In altricial avian species, for whom early-life sustenance is reliant on parental provisioning, intricate family dynamics emerge among individual chicks, their siblings, and their parents (e.g. Davies, 1976; Wright & Leonard, 2002). An increased brood size presents numerous challenges for parents and offspring. For parents, having to care for more offspring increases care demands, forcing parents to either reallocate resources that would otherwise be used for their own somatic maintenance (compensation hypothesis) or increase their own energy intake (increased-intake hypothesis) (Nilsson, 2002). For nestlings, being raised in larger broods results in higher rates of sibling competition for parental resources (Godfray, 1995; Godfray & Parker, 1992; Grodzinski & Johnstone, 2012) and the prioritization of investment in the development of structures that enhance their competitive ability, such as their gapes (Gil et al., 2008). Ultimately, being raised in a larger brood can not only lead to reduced growth rates and nestling mortality but also have long-term effects on fitness prospects in adulthood (Boonekamp et al., 2014, Boonekamp et al., 2020; Briga et al., 2017; Dijkstra et al., 1990; Hõrak, 2003; Nicolaus et al., 2009).

Begging is the main behavior displayed by the chicks in the nest to attract the parents’ attention and compete for parental resources with siblings (Godfray & Johnstone, 2000). Begging consists of intense and conspicuous vocalizations as well as head and body movements (e.g. wing flapping, gaping, Kilner & Johnstone, 1997; Zann, 1996). Theoretical work on begging behavior often relies on the assumption that begging entails a cost, either physiological as increased energy expenditure or by increasing predation risk. Experimental studies have confirmed that nestling begging calls can increase the risk of predation (Haskell, 1994, 1999; Leech & Leonard, 1997). The energetic costs of begging appear modest (Leech & Leonard, 1996; Mccarty, 1996), but even the observed energetic costs may constitute a substantial part of the energy that can be otherwise allocated to growth (Verhulst & Wiersma, 1997) as confirmed by experiments that manipulated begging effort independent from the amount of food received (Nettle et al., 2017; Rodriguez-Girones et al., 2001; Soler et al., 2014).

Given that begging is a possibly dangerous and energetically costly behavior, theory has been developed to understand how this behavior has evolved. Central in theoretical considerations is that there is an inherent evolutionary conflict between parents and chicks: due to divergent genetic interests, the optimal level of parental care from the perspective of the offspring is usually higher than the optimal level from the perspective of the parents, hence there is a “parent-offspring conflict” (Trivers, 1974). In this context, begging may serve as an honest signal of need, enabling parents to make informed decisions on the allocation of resources to the brood and between nestlings (Godfray, 1991; Royle et al., 2002). Offspring need, and possibly thereby the optimal begging effort, depends on brood size, because brood size manipulation typically modulates nestling growth (Burness et al., 2000; de Kogel, 1997). At the same time, competition for food is likely to increase with increasing brood size unless parents increase provisioning rate sufficiently to fully match the demands of an increased brood size. Both the increased level of competition and incomplete parental compensation for a larger brood are likely to increase ‘nestling need’, and hence begging effort (Harper, 1986).

In zebra finches (*Taeniopygia castanotis*), it is well established that being raised in larger broods has multiple phenotypic effects that reduce fitness prospects, possibly through reduced growth (Briga et al., 2017; de Kogel, 1997; Griffith & Buchanan, 2010). These effects include reduced immunocompetence (Naguib et al., 2004), higher standard metabolic rate (Verhulst et al., 2006, but see Briga & Verhulst, 2021), and altered mating preferences (Holveck & Riebel, 2010). Our aim was to determine whether diminished growth results from increased resource allocation to competition, a *per capita* reduction in parental provisioning rate, or a combination of the two. We therefore cross-fostered zebra finch chicks into broods of either two or six young and filmed the manipulated broods to record the behavior of parents and offspring. We envisaged different scenarios that could emerge, depending on responses of parents and offspring to the brood size manipulation. If provisioning behavior and begging effort remain unchanged, the resulting reduction in *per capita* provisioning would depress growth. If parents maintain the same *per capita* provisioning rate, while the offspring increase their begging effort, growth in large broods would be reduced due to increased resource allocation to begging. Intermediate responses of both parents and offspring would lead to reduced growth due to a combination of reduced *per capita* provisioning and increased resource allocation to begging.

## Methods

### Cross fostering

Zebra finch pairs were randomly matched and placed in breeding cages (L x H x D: 80 × 40 × 40 cm) on a 14L:10D schedule at ~ 25^°^C temperature and ~60% humidity. In the cages, there were nestboxes, nesting material (hay) as well as food and water *ad libitum*. Up until the hatching of the first egg in the nest, egg food was also supplied to the parents. Nestboxes were checked daily for the presence of eggs or chicks. Chicks were cross-fostered until the maximum age of five days to broods of either two (n=21) or six (n=14) chicks, which matches the brood size range observed in the wild (Zann, 1996). On the day of cross-fostering, all chicks were weighed and marked by clipping one or more of the head tufts for individual recognition. Captive zebra finch eggs hatch asynchronously, and age differences within broods remained similar after manipulation. The mean age at cross-fostering was 3.2±1.5 days. No siblings were placed in the same nests and no parents were taking care of their own offspring. Chicks were ringed at the age of 12 days, and weighed again at 15 and 35 days. At 35 days, having reached nutritional independence (Zann, 1996), chicks were moved from the breeding cage into larger indoor aviaries with two pairs of foster parents for sexual imprinting.

Extra-pair fertilizations are rare in natural zebra finch populations (Griffith et al., 2010; Zann, 1996), and we assume therefore that as a rule the species has evolved to perceive other brood members as siblings. This is relevant insofar that potential effects of variation in relatedness within the families on interactions between brood members and parents can be ignored in the context of this study.

### Camera observations

Cameras were placed above the nest an hour before the start of the recording, to allow the birds to adjust to them. Each observation consisted of an hour of filming per nest and each nest was recorded multiple times at multiple ages of the chicks until they fledged. The video recordings were then uploaded and analyzed in BORIS, an interactive software program for behavioral observations (Friard & Gamba, 2016). Frequency and duration of the following behaviors of chicks and parents was recorded:

- *Resting*, the individual rests and does nothing else or the adult is just sitting in the nest;
- *Begging*, the chick makes mouth and body movements to extort parental resources;
- *Feeding*, the chick takes food from the parent or the parent gives food to the chick (foraging time not included for the parents).
- *Cleaning*, the individual (either chick or parent) cleans its feathers;
- *Moving*, the individual (either chick or parent) moves within the nest or the parent makes changes in the nest (e.g., nest-building);
- *Other*, the individual does not display any of the behaviors mentioned above, is outside of the nest or is not visible.

We recorded 443 individual observations of chicks and 240 of adults in / rearing small broods and 634 observations of chicks and 100 of adults in / rearing large broods. Two chicks were observed in each video, randomly selected. Note that we could track individuals for the duration of the video but could not identify the chicks individually. The distribution of ages at which the chick observations were made can be found in Figure S1.

### Statistical analyses

All analyses were performed in R v.4.1.2 (R Core Team, 2023). All models were fitted using package *lme4* v. 1.1.35.1 (Bates et al., 2015) and we tested the statistical significance of variables and their interactions using the package *car* v. 3.1.2 (Fox & Weisberg, 2019), and visually inspected the residuals to verify adherence to the model assumptions. When analyzing variation in proportion of time, proportions were arcsine square root transformed (θ= arcsin √p, where p is the proportion) prior to analysis to meet the homoscedasticity assumption. Video recordings were made in different breeding rounds and analyzed by different observers (n=3), and to account for possible differences between breeding rounds and the co-varying identity of the observers (that may differ in the details of scoring behavior) we included observed ID as random effect in the analyses. Video recordings were made at different ages, and we grouped observations in age categories (0-4, 5-9, 10-5 and 16+ days) to avoid losing too many degrees of freedom when chick age effects on provisioning.

### Chick mass

We used linear mixed models to determine the effect of brood size on chick mass. To account for variations in the age at cross-fostering, we included the age at which mass was recorded as a covariate. Since not all chicks were cross-fostered and therefore weighed at the same age, the average age at cross-fostering per brood served as the reference point for the first timepoint, and day 15 served as the second timepoint for each observation. As random effects we included identity of each chick and brood. We used the F-statistic metric derived from a Type III Analysis of Variance Table with Satterthwaite’s method to test the significance of the interactions between brood size and age category. Note that mass data was not available for all chicks included in the experiment.

### Chick behavior

We used linear mixed models with the arcsine transformed proportion of time spent on each behavior per observation as the response variable using brood size and age category at recording as explanatory variables while brood and observer identities and were included as random intercepts.

### Parent behavior

We used linear mixed models of time spent on feeding per parent as the response variable using brood size as the explanatory variable and included brood identity as random effect. Since parental behavior was not always recorded, parent provision rate to the broods were calculated from chick provision rate data by extrapolating the observed provisioning rate from the observed chicks to the entire brood. Specifically, because chicks were observed in each video, we averaged the proportion of time the two chicks were observed being fed per observation. In order to account for chick mortality within the brood, we then multiplied by the number of chicks still alive in the nest at the time of observation (usually two or six as mortality was very low). Lastly, we divided the result by two to obtain the proportion of time spent feeding per parent (sex of the feeding parent was not always recorded).

## Results

### Chick mass

Brood size affected chick growth as evidenced by a significant interaction between age and brood size, confirming diminished growth rate in chicks in large broods (Table 1). Being raised in large broods resulted in gaining 0.04±0.2 g less per day (Fig.1a). Age at cross-fostering did not significantly increase the explained variance (*p*=0.80) and was therefore excluded from the final model. At age 15 days, chicks raised in large broods weighed on average 0.5±2.33 g less than chick reared in small broods (Fig.1b).

**Table 1.** Linear mixed effects model results of the effects of brood size and age on chick mass (gr) (N=110 observations, from 20 small broods and 13 large broods).

|  | Chick mass (gr) |  |  |  |
| --- | --- | --- | --- | --- |
| | Estimate (± SE) | t-value | df | Pr(>\|t\|) |
| *Fixed effects* |  |  |  |  |
| Intercept | 0.34 (± 0.30) | 1.15 | 77.92 | 0.25 |
| large brood | 0.13 (± 0.45) | 0.3 | 63.32 | 0.77 |
| age (days) | 0.75(± 0.02) | 37.09 | 98.14 | < 0.001 |
| large brood * age | -0.07 (± 0.03) | -2.29 | 98.68 | 0.02 |
| *Random effects* |  |  |  |  |
| bird id | 0.12 |  |  |  |
| brood id | 0.7 |  |  |  |
| Residual | 1.33 |  |  |  |

**Figure 1.**
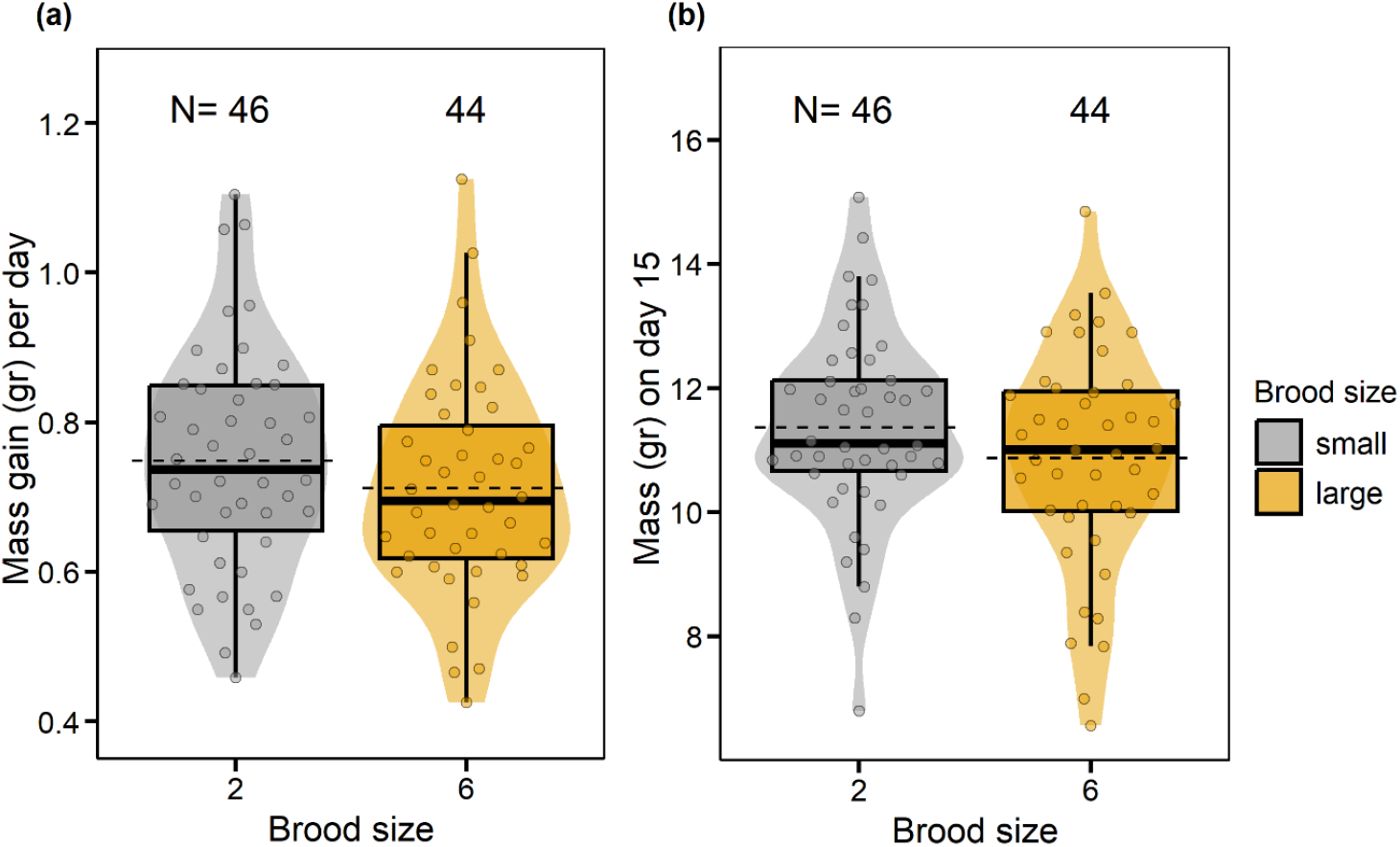
Mass gain per day of chicks raised in small and large broods (a). Mass of chicks raised in small and large broods on day 15 (b). Violin plot widths represent the data density, boxplots show the median and quartiles while dashed lines show the average daily mass gain and average mass on day 15 for chicks raised in small and large broods.

### Begging behavior

Chicks in large broods begged on average 7% more than chicks from small broods. This increase was not at the expense of resting time, which was not affected by brood size, but rather of other behaviors (Fig.2). The proportion of time at which chicks displayed each behavior per age category is summarized in Figure S2. Chicks in large broods consistently begged more throughout the nestling period, independent of age (F_1,166_=0.06, *p*<0.05). Begging intensity peaking between day five and fifteen (Table 2, Fig.3).

**Table 2.** Linear mixed effects model results brood size and age effects on the proportion of time (arcsine square root transformed) that nestlings spent begging (N=195 observations, from 21 small broods and 14 large broods). Note that observed identity is fully confounded with breeding round and effects of breeding round and observer identity can therefore not be separated.

|  | Proportion of time chicks spent begging |  |  |  |
| --- | --- | --- | --- | --- |
| | Estimate (± SE) | t-value | df | Pr(>\|t\|) |
| *Fixed effects* |  |  |  |  |
| Intercept | 0.0089 ± 0.0041 | 2.2 | 4.98 | 0.08 |
| large brood | 0.012 ± 0.0022 | 5.71 | 23.28 | <0.001 |
| age 5-9 days | 0.017 ± 0.0027 | 6.3 | 189 | <0.001 |
| age 10-15 days | 0.014 ± 0.0028 | 5.2 | 171.2 | <0.001 |
| age 16+ days | 0.0028 ± 0.0031 | 0.89 | 191.7 | 0.39 |
| *Random effects* |  |  |  |  |
| brood id | 1.59e-05 |  |  |  |
| observer | 2.72e-05 |  |  |  |
| residual | 1.08e-04 |  |  |  |

**Figure 2.**
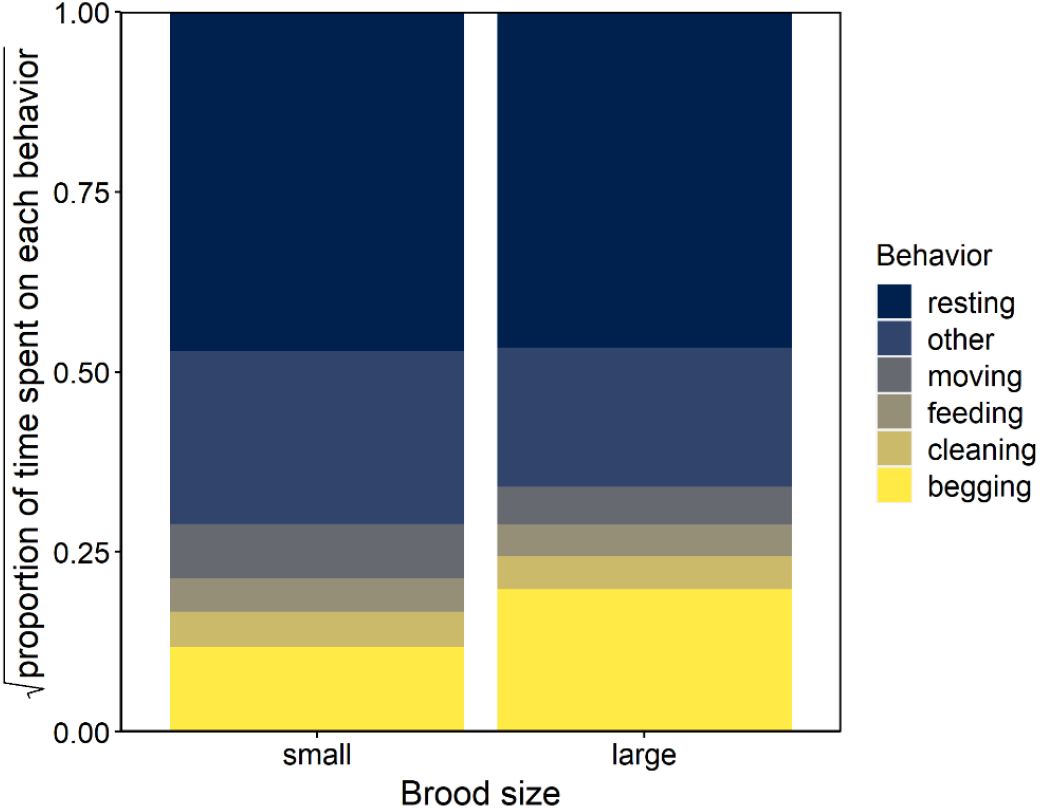
Proportion of time chicks in small and large broods spent on each behavior over all ages pooled. Proportions of each behavior per observation were normalized and then square root transformed. See Figure S1 for the results for the different age categories.

**Figure 3.**
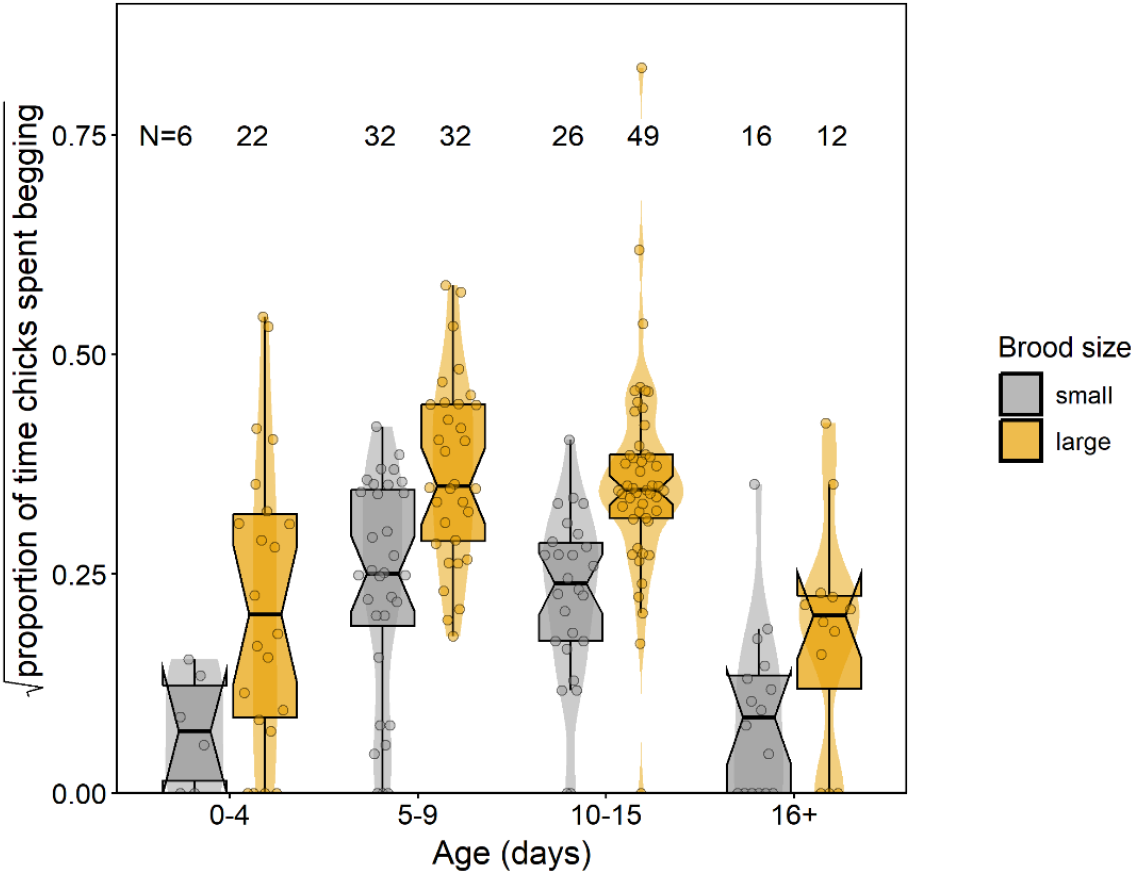
Proportion of time chicks in small and large broods spent begging over the nestling period (N=195 observations). For graphical purposes proportions were square root transformed. Violin plot widths represent the data density, boxplots show the median and quartiles.

### Parent provisioning

Parents caring for large broods spent 7.7% and parents caring for small broods spent 0.9% of their time transferring food to the chicks (Fig.4, Table 3). The amount of time parents spent feeding their brood did not significantly differ between the early days of the chicks’ lives and the later age categories (Fig.4, Table 3).

**Table 3.**
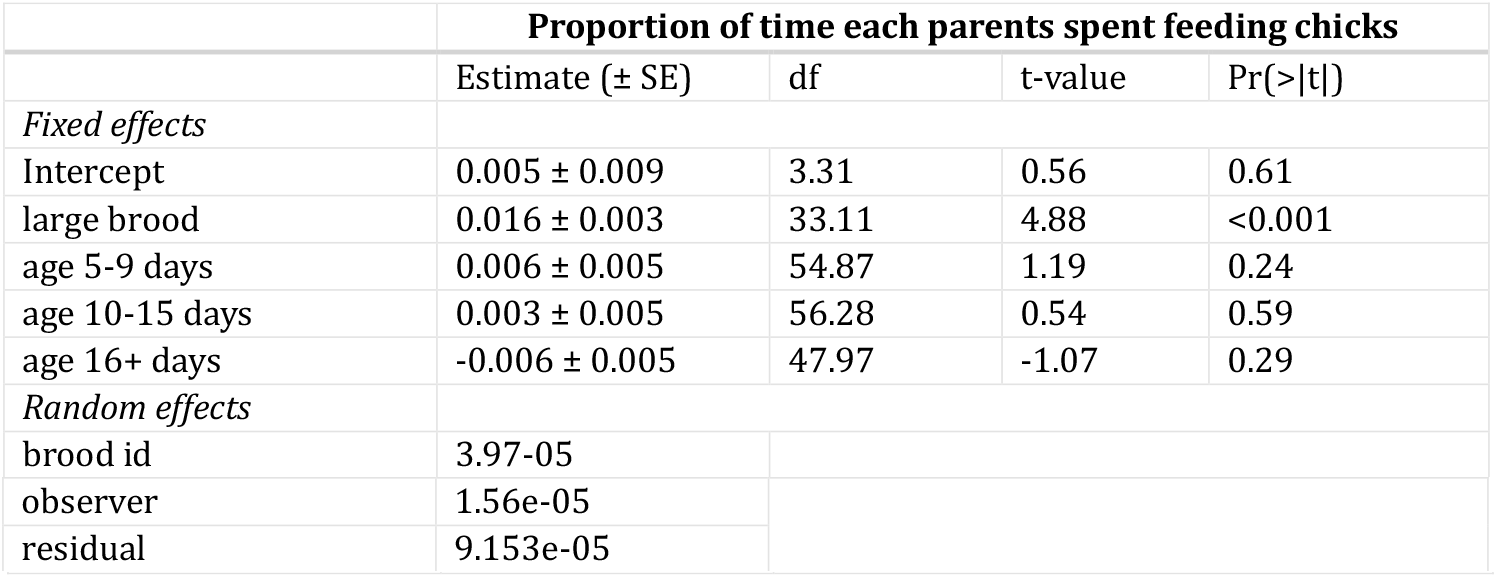
Linear mixed effects model results of brood size effects on the proportion of time (arcsine square root transformed) parents spent feeding their chicks (N=71 observations, from 21 small broods and 14 large broods).

|  | Proportion of time each parents spent feeding chicks |  |  |  |
| --- | --- | --- | --- | --- |
| | Estimate (± SE) | df | t-value | Pr(>\|t\|) |
| *Fixed effects* |  |  |  |  |
| Intercept | 0.005 ± 0.009 | 3.31 | 0.56 | 0.61 |
| large brood | 0.016 ± 0.003 | 33.11 | 4.88 | <0.001 |
| age 5-9 days | 0.006 ± 0.005 | 54.87 | 1.19 | 0.24 |
| age 10-15 days | 0.003 ± 0.005 | 56.28 | 0.54 | 0.59 |
| age 16+ days | -0.006 ± 0.005 | 47.97 | -1.07 | 0.29 |
| *Random effects* |  |  |  |  |
| brood id | 3.97-05 |  |  |  |

|  |  |
| --- | --- |
| observer | 1.56e-05 |
| residual | 9.153e-05 |

**Figure 4.**
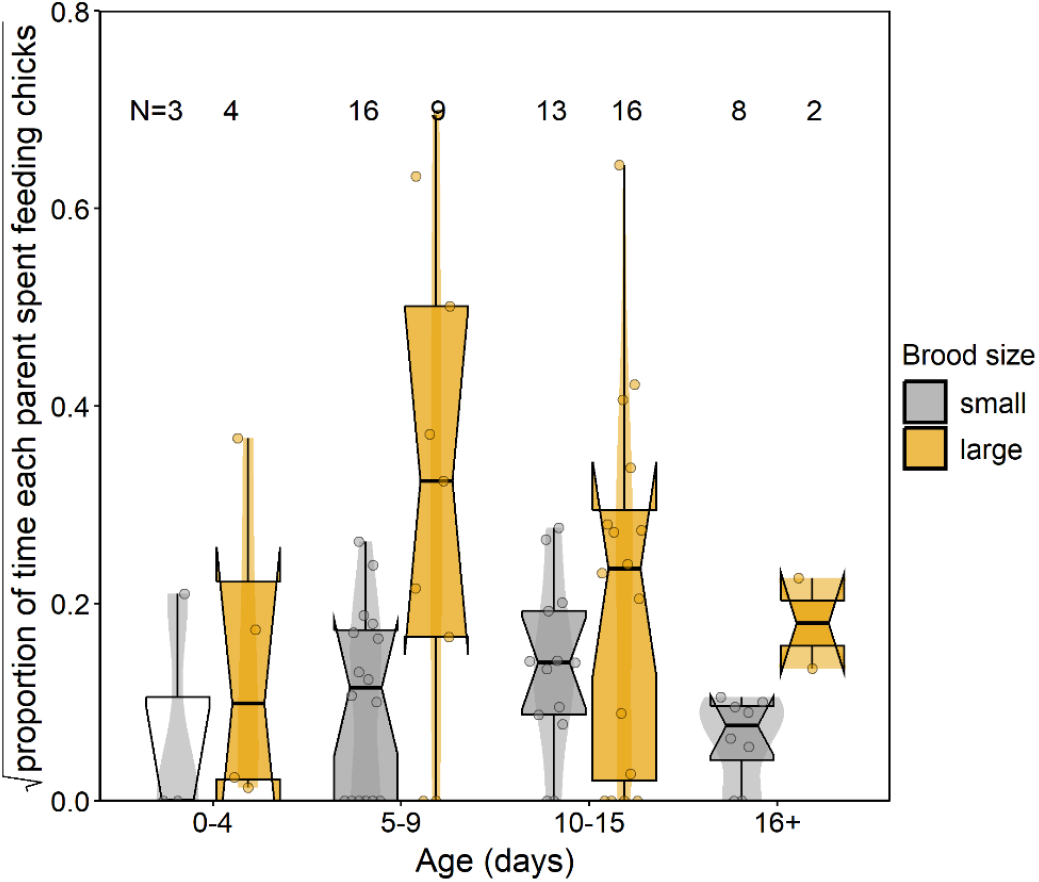
Proportion of time each parent spent transferring food to the chicks in small and large broods over the nestling period (N=71 observations). For graphical purposes proportions were square root transformed. Violin plot widths represent the data density, boxplots show the median and quartiles.

### Per capita chick feeding

Chicks reared in large broods spent 0.16% more time being fed than chicks in small broods (Fig.5, Table 4). The interaction between brood size and age category did not explain a significant part of the variance when added to the model (F_1,179_=0.81, *p*>0.05). The proportion of time being fed peaked between five and fifteen days for chicks of both brood sizes (Fig.5b).

**Table 4.** Linear mixed effects model results of brood size and age effects on proportion of time (arcsine square root transformed) individual chicks spent being fed (N=194 observations, from 21 small broods and 14 large broods).

|  | Proportion of time chicks spent being fed |  |  |  |
| --- | --- | --- | --- | --- |
| | Estimate (± SE) | df | t-value | Pr(>\|t\|) |
| *Fixed effects* |  |  |  |  |
| Intercept | 0.0037 ± 0.0039 | 2.33 | 0.96 | 0.42 |
| large brood | 0.0039 ± 0.0013 | 29.95 | 2.88 | <0.01 |
| age 5-9 days | 0.0029 ± 0.0012 | 184.37 | 2.48 | 0.014 |
| age 10-15 days | 0.0016 ± 0.0012 | 188.05 | 1.29 | 0.20 |
| age 16+ days | -0.0029 ± 0.0014 | 180.41 | -2.11 | 0.037 |
| *Random effects* |  |  |  |  |
| brood id | 1.24e-05 |  |  |  |
| observer | 3.89e-05 |  |  |  |
| residual | 1.863e-05 |  |  |  |

**Figure 5.**
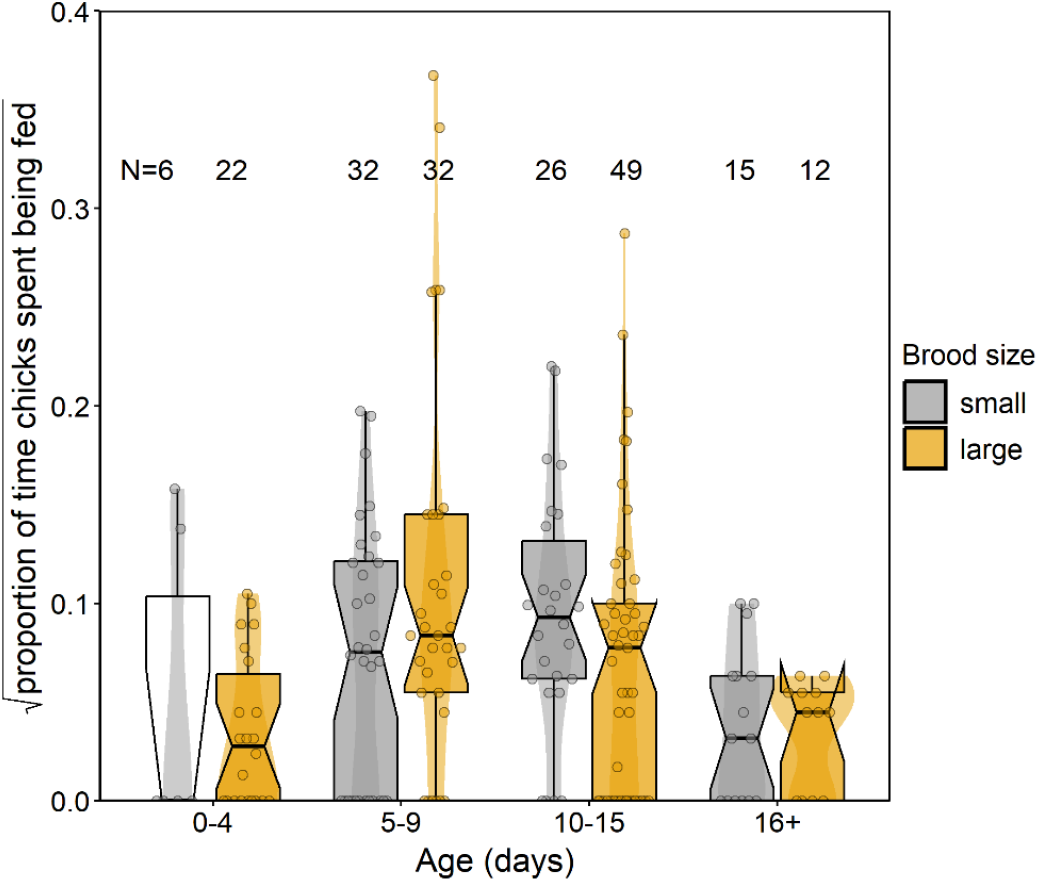
Proportion of time chicks in small and large broods spent being fed over the nestling period (N=194 observations). For graphical purposes proportions were square root transformed. Violin plot widths represent the data density, boxplots show the median and quartiles.

## Discussion

It is well established that being raised in a larger brood adversely affects growth – even in captive zebra finches with an *ad libitum* food supply. To shed light on the mechanistic underpinnings of this effect we manipulated brood size and recorded parent and offspring behavior throughout the nestling period. We found that zebra finch nestlings raised in large broods are faced with a diminished growth rate, despite a slight increase in *per capita* feeding rate when compared to chicks raised in small broods. We hypothesize that the diminished growth results from increased resource allocation to competition through begging, which is consistent with earlier studies across various bird species, which have shown that nestlings in larger broods beg more intensively (Leonard et al., 2000; Neuenschwander et al., 2003; Wright et al., 2002) as well as empirical studies that demonstrate begging to have energetic, metabolic and other physiological costs (Abraham & Evans, 1999; Bachman & Chappell, 1998; Roulin, 1998). Our findings suggest a complex trade-off between competitive behavior for survival and physical development in the nestling period of zebra finches, also supporting pervious claims about a significant energetic cost associated with begging.

When parents have to care for more chicks, they can adaptively adjust their provisioning rate to match the needs of a larger brood as displayed by the increased parental feeding for parents and per capita provisioning rates for chicks in large broods. Previous research has demonstrated that larger broods are generally associated with increased parental provisioning rates in birds (Bowers et al., 2014; Lessells, 1993; Stoehr et al., 2001). Parents caring for large broods in our study spent more time feeding their chicks. This increased provisioning is likely, at least in part, a response to the increase in begging, given that begging playback experiments successfully elevated parental provisioning (e.g. Burford et al., 1998; Ottosson et al., 1997; Price, 1998; Tarwater et al., 2009). In most studies, the effect of brood size manipulation on parental provisioning only partly compensated for the change in brood size, and, as a consequence, *per capita* provisioning rate is typically lower in enlarged broods (e.g. Dijkstra et al., 1990). Our study is unusual in this respect, in that the brood size effect on provisioning rate was such that there was an *increase* in *per capita* provisioning rate in large broods. Begging is the foraging technique of altricial nestlings, and, as any foraging technique, the yield will depend on food availability in the environment. In our study, food availability was *ad libitum* from the perspective of the parents, which is likely to have shaped our results, in that the energy yield of begging was probably higher than it would have been in a more natural environment where parents usually have to make a greater effort to gather food.

With respect to the question how the brood size effect on nestling growth arises, we infer from the present findings that brood size increased *per capita* provisioning rate and that the brood size effect on growth must therefore be due to other factors than the amount of food received. Instead, we consider the increased energy allocation to begging the most likely cause of the brood size effect on nestling growth. Our findings thereby provide indirect evidence for a significant energetic cost associated with begging behavior, similar to experiments in which nestlings were forced to beg at a higher rate for the same amount of food, which also depressed growth (Nettle et al., 2017; Rodriguez-Girones et al., 2001; Soler et al., 2014). A new dimension revealed by the present study is that family dynamics can induce a qualitatively similar effect on growth in the absence of an experimentally set limit on how much food the nestlings consume. Interestingly, this suggests that the brood size effect on growth is in a way self-inflicted, because when parents had adjusted their provisioning rate spontaneously, i.e., without having been stimulated to do so by the nestling’s begging intensity and assuming that the parents feed chicks in the same rate during feeding, nestlings in small and large broods could have allocated the same amount of energy to growth. Paying the begging costs, which affects inclusive fitness of all family members, may be the evolutionary stable solution that is the outcome of the parent-offspring conflict over the optimal level of parental care.

## Acknowledgements

We thank the animal caretakers of the University of Groningen as well as numerous students including Gillian Bakx and Wietske van Buiten for assistance with data collection. MB thanks the funding from the Turku Collegium for Science, Medicine and Technology.

## Ethics approval

All methods and experiments for the zebra finches detailed in this manuscript were performed under approval of the Central Committee for Animal Experiments (Centrale Commissie Dierproeven) of the Netherlands, under license AVD1050020174344.

## Data availability statement

The data and scripts can be found in https://github.com/tangilim/ZF_Begging.

## Supplementary information

**Figure S1.**
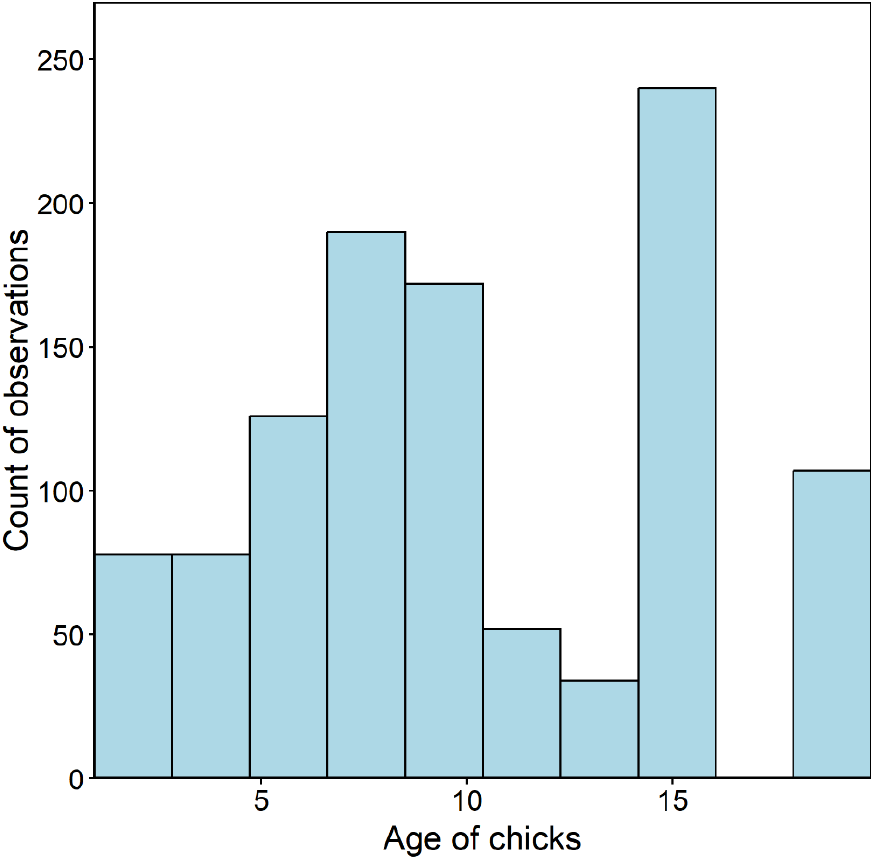
Age of the chicks when observations on chick behavior were recorded (N=1077 observations).

**Figure S2.**
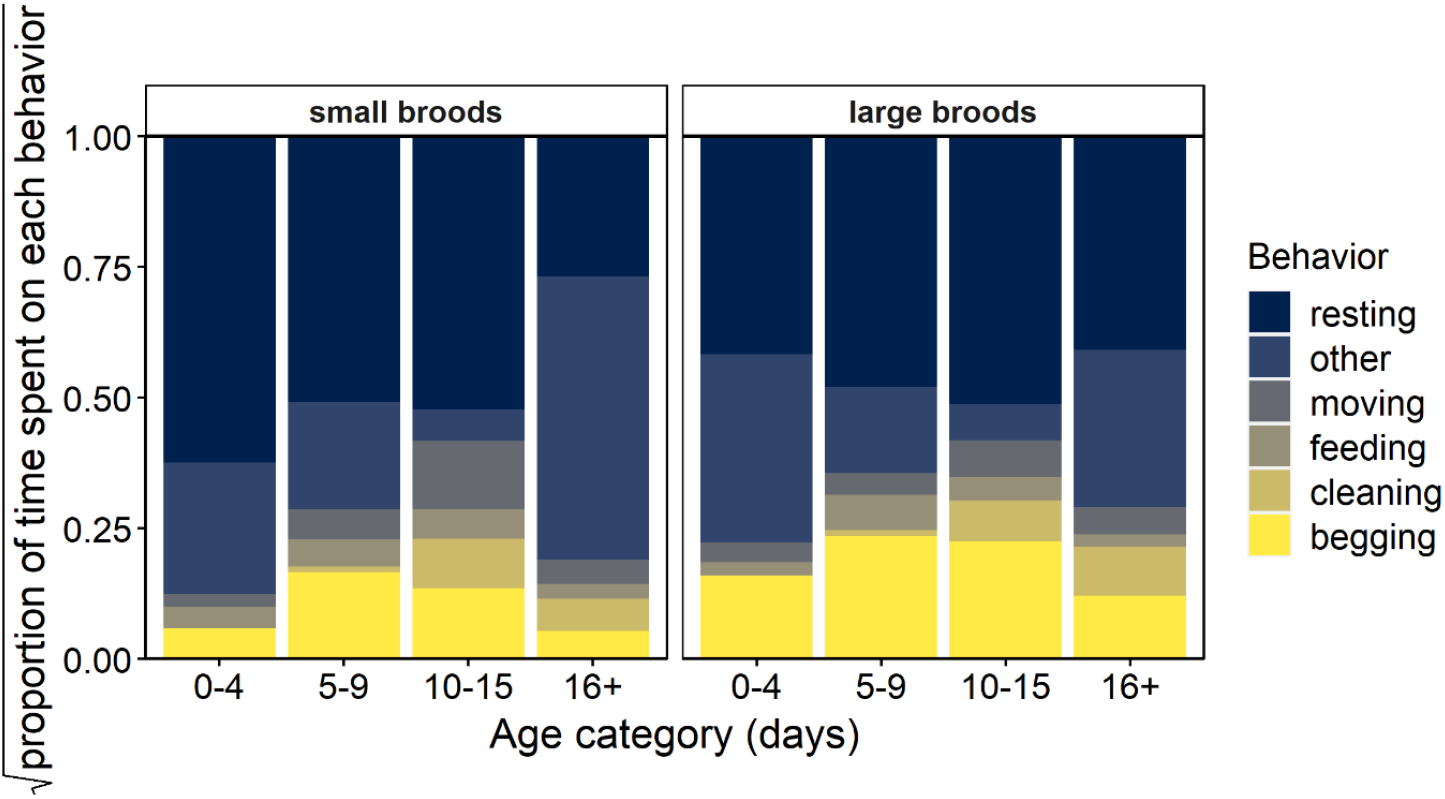
Proportion of time spent on each behavior by chicks in small and large broods divided in the age categories of the chicks. For graphical purposes proportions of each behavior per observation were normalized and then square root transformed.

## References

Abraham, C. L., & Evans, R. M. (1999). Metabolic costs of heat solicitation calls in relation to thermal need in embryos of American white pelicans. Animal Behaviour, 57, 967–975. 10.1006/anbe.1998.1060

Andrews, C., Viviani, J., Egan, E., Bedford, T., Brilot, B., Nettle, D., & Bateson, M. (2015). Early life adversity increases foraging and information gathering in European starlings, Sturnus vulgaris. Animal Behaviour, 109(109), 123–132. 10.1016/j.anbehav.2015.08.009

Bachman, G. C., & Chappell, M. A. (1998). The energetic cost of begging behaviour in nestling house wrens. Animal Behaviour, 55, 1607–1618. 10.1006/anbe.1997.0719

Bates, D., Ma chler, M., Bolker, B., & Walker, S. (2015). Fitting Linear Mixed-Effects Models Using {lme4}. Journal of Statistical Software, 67(1), 1–48. 10.18637/jss.v067.i01

Boonekamp, J. J., Bauch, C., & Verhulst, S. (2020). Experimentally increased brood size accelerates actuarial senescence and increases subsequent reproductive effort in a wild bird population. Journal of Animal Ecology, 89(6), 1395–1407. 10.1111/1365-2656.13186

Boonekamp, J. J., Mulder, G. A., Salomons, H. M., Dijkstra, C., & Verhulst, S. (2014). Nestling telomere shortening, but not telomere length, reflects developmental stress and predicts survival in wild birds. Proceedings of the Royal Society B: Biological Sciences, 281(1785), 1– 7. 10.1098/rspb.2013.3287

Bowers, E. K., Nietz, D., Charles, F., & Sakaluk, S. K. (2014). Parental provisioning in house wrens : effects of varying brood size and consequences for offspring. Behavioral Ecology, 25(6), 1485–1493. 10.1093/beheco/aru153

Briga, M., Koetsier, E., Boonekamp, J. J., Jimeno, B., & Verhulst, S. (2017). Food availability affects adult survival trajectories depending on early developmental conditions. Proceedings of the Royal Society B: Biological Sciences, 284(1846), 20162287. 10.1098/rspb.2016.2287

Briga, M., & Verhulst, S. (2021). Mosaic metabolic ageing: Basal and standard metabolic rates age in opposite directions and independent of environmental quality, sex and life span in a passerine. Functional Ecology, 35(5), 1055–1068. 10.1111/1365-2435.13785

Burford, J. E., Friedrich, T. J., & Yasukawa, K. (1998). Response to playback of nestling begging in the red-winged blackbird, Agelaius phoeniceus. Animal Behaviour, 56(3), 555–561. 10.1006/anbe.1998.0830

Burness, G. P., McClelland, G. B., Wardrop, S. L., & Hochachka, P. W. (2000). Effect of brood size manipulation on offspring physiology: An experiment with passerine birds. Journal of Experimental Biology, 203(22), 3513–3520. 10.1242/jeb.203.22.3513

Davies, N. B. (1976). Parental Care and the Transition To Independent Feeding in the Young Spotted Flycatcher (Muscicapa Striata). Behavior, 280–294.

de Kogel, C. H. (1997). Long-Term Effects of Brood Size Manipulation on Morphological Development and Sex-Specific Mortality of Offspring. The Journal of Animal Ecology, 66(2), 167. 10.2307/6019

Dijkstra, C., Bult, A. S.B.,, Daan, S., Meijer, T., & ZIilster, M. (1990). Brood Size Manipulations in the Kestrel (Falco tinnunculus): Effects on Offspring and Parent Survival. Journal of Animal Ecology, 59(1), 269–285.

Fox, J., & Weisberg, S. (2019). An {R} Companion to Applied Regression (Third). Sage. https://socialsciences.mcmaster.ca/jfox/Books/Companion/

Friard, O., & Gamba, M. (2016). BORIS : a free, versatile open-source event-logging software for video / audio coding and live observations. Methods in Ecology and Evolution, 7, 1325–1330. 10.1111/2041-210X.12584

Gerritsma, Y. H. (2025). From adversity to personality. The effect of early-life adversity on zebra finch behaviour.

Gil, D., Bulmer, E., Celis, P., & Lo pez-Rull, I. (2008). Adaptive developmental plasticity in growing nestlings: Sibling competition induces differential gape growth. Proceedings of the Royal Society B: Biological Sciences, 275(1634), 549–554. 10.1098/rspb.2007.1360

Godfray, H. C. J. (1991). Signaling of need by offspring to their parents. Nature, 352, 328–330.

Godfray, H. C. J. (1995). Signaling of need between parents and young: parent-offspring conflict and sibling rivalry. The American Naturalist, 146(1), 1–24.

Godfray, H. C. J., & Johnstone, R. (2000). Begging and bleating : the evolution of parent − offspring signalling Begging and bleating : the evolution of parent – offspring signalling. Philosophical Transactions of the Royal Society B: Biological Sciences, 355, 1581–1591. 10.1098/rstb.2000.0719

Godfray, H. C. J., & Parker, G. (1992). Sibling competition, parent-offspring conflict and clutch size. Animal Behaviour, 43(1974), 473–490.

Goldenberg, S. Z., & Wittemyer, G. (2018). Orphaning and natal group dispersal are associated with social costs in female elephants. Animal Behaviour, 143, 1–8. 10.1016/j.anbehav.2018.07.002

Griffith, S. C., & Buchanan, K. L. (2010). Maternal effects in the Zebra Finch: A model mother reviewed. Emu, 110(3), 251–267. 10.1071/MU10006

Griffith, S. C., Holleley, C. E., Mariette, M. M., Pryke, S. R., & Svedin, N. (2010). Low level of extrapair parentage in wild zebra finches. Animal Behaviour, 79(2), 261–264. 10.1016/j.anbehav.2009.11.031

Grodzinski, U., & Johnstone, R. A. (2012). Parents and offspring in an evolutionary game : the effect of supply on demand when costs of care vary. Proceedings of the Royal Society B: Biological Sciences, 279, 109–115. 10.1098/rspb.2011.0776

Harper, A. B. (1986). The evolution of begging: sibling competition and parent-offspring conflict. American Naturalist, 128(1), 99–114. 10.1086/284542

Haskell, D. G. (1994). Experimental evidence that nestling begging behaviour incurs a cost due to nest predation. Proceedings of the Royal Society B: Biological Sciences, 257(1349), 161–164. 10.1098/rspb.1994.0110

Haskell, D. G. (1999). The effect of predation on begging-call evolution in nestling wood warblers. Animal Behaviour, 57(4), 893–901. 10.1006/anbe.1998.1053

Holveck, M. J., & Riebel, K. (2010). Low-quality females prefer low-quality males when choosing a mate. Proceedings of the Royal Society B: Biological Sciences, 277(1678), 153–160. 10.1098/rspb.2009.1222

Hõrak, P. (2003). When to pay the cost of reproduction? A brood size manipulation experiment in great tits (Parus major). Behavioral Ecology and Sociobiology, 54(2), 105–112. 10.1007/s00265-003-0608-1

Kilner, R. M., & Johnstone, R. A. (1997). Begging the question: are offspring solicitation behaviors signals of need? Trends in Ecology & Evolution, 12(1), 11–15.

Leech, S. M., & Leonard, M. L. (1996). Is there an energetic cost to begging in nestling tree swallows (Tachycineta bicolor)? Proceedings of the Royal Society B: Biological Sciences, 263(1373), 983–987. 10.1098/rspb.1996.0145

Leech, S. M., & Leonard, M. L. (1997). Begging and the risk of predation in nestling birds. Behavioral Ecology, 8(6), 644–646. 10.1093/beheco/8.6.644

Leonard, M. L., Horn, A. G., Gozna, A., & Ramen, S. (2000). Brood size and begging intensity in nestling birds. Behavioral Ecology, 11(2), 196–201.

Lessells, C. M. (1993). The cost of reproduction: do experimental manipulations measure the edge of the options set. Etologia, 3, 95–111.

Mainwaring, M. C., Blount, J. D., & Hartley, I. A. N. R. (2012). Hatching asynchrony can have longterm consequences for offspring fitness in zebra finches under captive conditions. Biological Journal of the Linnean Society, 106, 430–438.

Mccarty, J. P. (1996). The energetic cost of begging in nestling passerines. The Auk, 113(1), 178– 188.

Mock, D. W., Schwagmeyer, P. L., & Dugas, M. B. (2009). Parental provisioning and nestling mortality in house sparrows. Animal Behaviour, 78(3), 677–684. 10.1016/j.anbehav.2009.05.032

Naguib, M., Riebel, K., Marzal, A., & Gil, D. (2004). Nestling immunocompetence and testosterone covary with brood size in a songbird. Proceedings of the Royal Society B: Biological Sciences, 271(1541), 833–838. 10.1098/rspb.2003.2673

Nettle, D., Andrews, C., Reichert, S., Bed, T., Kolenda, C., Parker, C., Martin-ruiz, C., Monaghan, P., & Bateson, M. (2017). Early-life adversity accelerates cellular ageing and affects adult inflammation : Experimental evidence from the European starling. Scientific Reports, 7(40794), 1–10. 10.1038/srep40794

Nettle, D., Monaghan, P., Gillespie, R., Brilot, B., Bedford, T., Bateson, M., & Nettle, D. (2015). An experimental demonstration that early-life competitive disadvantage accelerates telomere loss. Proceedings of the Royal Society B: Biological Sciences, 282(20141610), 1–8.

Neuenschwander, S., Brinkhof, M. W. G., & Ko, M. (2003). Brood size, sibling competition, and the cost of begging in great tits (Parus major). 14(4), 457–462.

Nicolaus, M., Michler, S. P. M., Ubels, R., Van Der Velde, M., Komdeur, J., Both, C., & Tinbergen, J. M. (2009). Sex-specific effects of altered competition on nestling growth and survival: An experimental manipulation of brood size and sex ratio. Journal of Animal Ecology, 78(2), 414–426. 10.1111/j.1365-2656.2008.01505.x

Nilsson, J.Å. (2002). Metabolic consequences of hard work. Proceedings of the Royal Society B: Biological Sciences, 269(1501), 1735–1739. 10.1098/rspb.2002.2071

Nilsson, J.Å., & Svensson, M. (2001). Sibling competition affects individual growth strategies in marsh tit, Parus palustris, nestlings. Animal Behaviour, 61, 357–365. 10.1006/anbe.2000.1602

Nussey, D. H., Kruuk, L. E. B., Morris, A., & Clutton-brock, T. H. (2007). Environmental conditions in early life influence ageing rates in a wild population of red deer. Current Biology, 17(23), 1000–1001.

Ottosson, U., Ba ckman, J., & Smith, H. G. (1997). Begging affects parental effort in the pied flycatcher, Ficedula hypoleuca. Behavioral Ecology and Sociobiology, 41(6), 381–384. 10.1007/s002650050399

Patterson, S. K., Strum, S. C., Silk, J. B., & Patterson, S. K. (2022). Early life adversity has long-term effects on sociality and interaction style in female baboons. Proceedings of the Royal Society B: Biological Sciences, 289(20212244), 1–9.

Price, K. (1998). Benefits of begging for yellow-headed blackbird nestlings. Animal Behaviour, 56(3), 571–577. 10.1006/anbe.1998.0832

R Core Team. (2023). R: A language and environment for statistical computing. (RStudio 2021.09.0+351 “Ghost Orchid” Release). https://www.r-project.org/

Rodriguez-Girones, M. A., Zuniga, J. M., & Redondo, T. (2001). Effects of begging on growth rates of nestling chicks. Behavioral Ecology, 12(3), 269–274.

Roulin, A. (1998). Forum On the cost of begging vocalization : implications of vigilance. Beha, 12(4), 506–515. 10.1093/BEHECO/12.4.506

Royle, N. J., Hartley, I. R., & Parker, G. A. (2002). Begging for control : when are offspring solicitation behaviours honest ? Trends in Ecology & Evolution, 17(9), 434–440.

Schwagmeyer, P., & Mock, D. (2008). Parental provisioning and offspring fitness : size matters. Animal Behaviour, 75, 291–298. 10.1016/j.anbehav.2007.05.023

Soler, M., Ruiz-Raya, F., Carra, L. G., Medina-Molina, E., Ibanez-Alamo, J.D., & Martín-Galvez, D. (2014). A long-term experimental study demonstrates the costs of begging that were not found over the short term. PLoS ONE, 9(11). 10.1371/journal.pone.0111929

Spagopoulou, F., Teplitsky, C., Lind, M. I., Chantepie, S., Gustafsson, L., & Maklakov, A. A. (2020). Silver-spoon upbringing improves early-life fitness but promotes reproductive ageing in a wild bird. Ecology Letters, 23(6), 994–1002. 10.1111/ele.13501

Stoehr, A. M., McGraw, K. J., Nolan, P. M., & Hill, G. E. (2001). Parental Care in Relation To Brood Size in the House Finch. Journal of Field Ornithology, 72(3), 412–418. 10.1648/0273-8570-72.3.412

Tarwater, C. E., Kelley, J. P., & Brawn, J. D. (2009). Parental response to elevated begging in a high predation, tropical environment. Animal Behaviour, 78(5), 1239–1245. 10.1016/j.anbehav.2009.07.040

Trivers, R. L. (1974). Parent-Offspring Conflict. American Zoologist, 264, 249–264.

Tung, J., Archie, E. A., Altmann, J., & Alberts, S. C. (2016). Cumulative early life adversity predicts longevity in wild baboons. Nature Communications, 7(11181), 1–7. 10.1038/ncomms11181

van de Pol, M., Bruinzeel, L. W., Heg, D., van der Jeugd, H., & Verhulst, S. (2006). A silver spoon for a golden future : long-term effects of natal origin on fitness prospects of oystercatchers (Haematopus ostralegus). Journal, 75, 616–626. 10.1111/j.13652656.2006.01079.x

Verhulst, S., Holveck, M. J., & Riebel, K. (2006). Long-term effects of manipulated natal brood size on metabolic rate in zebra finches. Biology Letters, 2(3), 478–480. 10.1098/rsbl.2006.0496

Verhulst, S., & Wiersma, P. (1997). Is begging cheap? The Auk, 114(1), 134.

Wright, J., Hinde, C., Fazey, I., & Both, C. (2002). Begging signals more than just short-term need: Cryptic effects of brood size in the pied flycatcher (Ficedula hypoleuca). Behavioral Ecology and Sociobiology, 52(1), 74–83. 10.1007/s00265-002-0478-y

Wright, J., & Leonard, M. L. (2002). The Evolution of Begging. Kluwer Academic Publishers.

Zann, R. A. (1996). The zebra finch. A synthesis of field and laboratory studies. Oxford University Press.

